# Age-Related Changes in Chimpanzee (*Pan troglodytes*) Cognition: Cross-Sectional and Longitudinal Analyses

**DOI:** 10.1101/2020.04.27.064626

**Authors:** William D Hopkins, Mary Catherine Mareno, Sarah J Neal Webb, Steve J Schapiro, Mary Ann Raghanti, Chet C Sherwood

## Abstract

Chimpanzees are the species most closely related to humans yet age-related changes in brain and cognition remain poorly understood. The lack of studies on age-related changes in cognition in chimpanzees is particularly unfortunate in light of the recent evidence demonstrating that this species naturally develops Alzheimer’s disease (AD) neuropathology. Here, we tested 213 young, middle-aged, and elderly chimpanzees on the Primate Cognitive Test Battery (PCTB), a set of 13 tasks that assess physical and social cognition in nonhuman primates. A subset of these chimpanzees (n=146) were tested a second time on a portion of the PCTB tasks as a means of evaluating longitudinal changes in cognition. Cross-sectional analyses revealed a significant quadratic association between age and cognition with younger and older chimpanzees performing more poorly than middle-aged individuals. Longitudinal analyses showed that, while young chimpanzees’ performance improved from test 1 to test 2, middle-aged and elderly chimpanzees’ performance showed significant decline over time. The collective data show that chimpanzees, like other nonhuman primates, show age-related decline in cognition. Further investigations into whether the observed cognitive decline is associated with AD pathologies in chimpanzees would be invaluable in understanding the comparative biology of aging and neuropathology in primates.

## Introduction

Decades of research in humans have documented age-related changes in cognition, brain and behavior (Abe et al., 2008; Bertrand et al., 2018; Peters, 2006). Some loss in cognitive functions and fidelity of cortical organization are considered a normal process of aging, but other decrements are attributable to specific neuropathological conditions, such as Alzheimer’s disease (AD) and frontotemporal dementia (FTD). Animal models, including primate models, have been developed as a means of understanding the mechanisms that underlie age-related changes in brain and cognition as well as facilitating the development of effective therapeutic interventions (Didier et al., 2016; Lacreuse & Herndon, 2008; McQuail & Nicolle, 2012; Phillips et al., 2014; Rosen et al., 2016; Rosen et al., 2011; Verdier et al., 2015; Walker & Jucker, 2017).

One species that is often overlooked in studies on age-related changes in cognition and the brain is chimpanzees (*Pan troglodytes*) (and other great apes). This is unfortunate given their genetic and neuroanatomical similarities to humans (Consortium, 2005; Sherwood et al., 2012) and more sophisticated cognitive and motor abilities (Tomasello & Call, 1997) when compared to more distantly related primates. Neuroimaging studies have reported small to moderate reductions in cortical surface area, gray matter thickness, gray matter volume and white matter connectivity in aged chimpanzees (Autrey et al., 2014; Chen et al., 2013; Raz et al., in press; Sherwood et al., 2011). Moreover, recent studies have documented the co-occurrence of both neurofibrillary tangles (NFTs) and β amyloid plaques (Aβ) in elderly chimpanzee postmortem brains (Edler et al., in press; Edler et al., 2017; Gearing et al., 1994; Rosen et al., 2008). The presence of both Aβ plaques and NFTs is the key neuropathological feature used to definitively confirm a diagnosis of AD in humans; thus, chimpanzees appear to be the only known species to naturally develop AD-like pathology. Additional studies involving chimpanzees could lead to important findings on the comparative biology of aging and may hold the key to understanding our own susceptibility to this neuropathological process (Edler et al., in press).

To date, there are only a handful of studies that have explicitly examined age-related changes in cognition in chimpanzees and all but one used cross-sectional rather than longitudinal designs which many have suggested are critically important in studies on aging (Raz et al., in press). In early work, Bernstein (1961) tested 8 young and 8 old chimpanzees and found no age differences in a series of object discrimination and on a wheel-rotating task. Riopelle and Rogers (1965) tested 19 chimpanzees (7 to 41 years of age divided into a young and old group) and reported no age-related deficits in discrimination tasks but did identify impairments in older chimpanzees on the shortest retention delays (0 and 5 s) of a delayed response and on an oddity task. In one recent study, Lacreuse et al. (2018) examined cognitive set switching in a modified version of the Wisconsin Card Sorting task in a sample of 30 female chimpanzees and found that older individuals made more perseveration errors and required more trials to reach criterion in the set switching dimensions than younger ones. Manrique and Call (2015) tested 25 apes (which included chimpanzees, bonobos, gorillas and orangutans) on a measure of cognitive flexibility and reported that younger and older individuals made significantly more perseveration errors compared to middle-aged apes.

In the lone longitudinal study in chimpanzees, Lacreuse et al. (2014) tested 38 females on the Primate Cognition Test Battery (PCTB), a 13-item test designed to assess social and physical cognition in nonhuman primates (Herrmann et al., 2007a; Herrmann, Hare, et al., 2010; Herrmann, Hernandez-Lloreda, et al., 2010; Joly et al., 2017; Russell et al., 2011; Schmitt et al., 2011). Lacreuse et al. (2014) found very little evidence of age-related decline in overall and physical cognition performance. In contrast, performance on measures of social cognition as well as motor function declined with age in this sample. Lacreuse et al. (2014) noted that the 4 oldest chimpanzees in the sample performed significantly worse over the 3-year study period than the remaining subjects on a measure of spatial memory.

In the current study, we tested for cross-sectional and longitudinal changes in cognition in a considerably larger sample of chimpanzees using the PCTB. From the cross-sectional data, we initially examined the linear and quadratic association between age and overall cognitive performance based on performance on the PCTB. We also tested a sub-sample of the chimpanzees three years later (on average) on a portion of the same PCTB tasks as a means of evaluating any longitudinal changes in performance with increasing age. We hypothesized that if chimpanzees exhibit age-related decline in cognitive functions, then both the cross-sectional and longitudinal data would reveal poorer performance in elderly individuals compared to middle-aged and young apes.

## Methods

### Ethics Statement

Investigation has been conducted in accordance with the ethical standards and according to the Declaration of Helsinki and according to national and international guidelines and has been approved by the authors’ institutional review board. All procedures involving the chimpanzees were approved by the local Institutional Animal Care and Use Committees, and adhered to the legal requirements of the United States and to the American Society of Primatologists’ Principles for the Ethical Treatment of Primates.

### Subjects

For the cross-sectional analysis, there were 218 chimpanzees (140 females, 78 males) ranging in age from 13 to 54 years (Mean 28.47, SD = 10.92) housed at two facilities, including the National Center for Chimpanzee Care (NCCC, n = 123), which is part of the MD Anderson Cancer Center, and Yerkes National Primate Research Center (YNPRC, n = 95) of Emory University. We classified the chimpanzees into three age groups including young (< 25 years, n = 98), middle-aged (25 to 36 years, n = 72) and elderly (> 37 years, n = 48). We selected these age cut-points based on life span and mortality data reported in Finch (2010) that indicate a sharp increase in chimpanzee mortality between 35 and 40 years of age. For the longitudinal analysis, the sample was comprised of 146 chimpanzees including 99 females and 47 males including 76 young, 46 middle-aged, and 24 elderly individuals. The range in time between the administration of the first and second PCTB tests varied from 1 to 7 years (Mean = 3.41, SD = 1.73).

### Procedures

Subjects were tested on a modified version of the primate cognition test battery (PCTB) originally described by Hermann et al. (Herrmann et al., 2007b; Herrmann, Hare, et al., 2010) and details have been described elsewhere (Russell et al., 2011). The PCTB attempts to assess subjects’ abilities in various areas of physical and social cognition. For our study, some aspects of the original PCTB were eliminated due to time and housing constraints. The previously published procedures were followed as similarly as possible but some tasks were modified to better address the questions at hand given the past experience and environmental constraints of our subjects. Each of the nine physical cognition tasks and the three social cognition tasks are described briefly below with notes made when procedures were altered from those described by Herrmann et al. (2007a). Subjects were generally tested in the order that the tasks are presented below and testing was completed over 1 to 5 testing sessions, depending on the motivation and attention of the subject.

### Physical Cognition Tasks

Nine tasks were utilized in the “Physical Cognition” portion of our test battery, including tasks exploring the apes’ spatial memory and understanding of spatial relationships, ability to differentiate between quantities, understanding of causality in the visual and auditory domains and their understanding of tools. Our test differed from the original PCTB in several ways. We excluded the Addition task as well as certain components of the Tool Properties tasks.

#### Spatial Memory (3 trials)

This test assessed subjects’ ability to remember the locations of food rewards. In this task, the subject watched as food was hidden in two of three possible locations. Each subject received all three possible combinations of baited locations. The subject was then allowed to search the locations. The subject was scored as successful if he/she located both food items without searching in the unbaited location.

#### Object Permanence (9 trials)

Here we tested an individuals’ ability to follow a food reward after invisible displacement given three different possible displacements. During single displacement trials, only one of three possible locations was manipulated and thus potentially baited. In the double displacement trials, two of three possible locations were manipulated, meaning that two of the three locations could potentially be baited. Double displacement trials were further divided by whether or not the baited locations were adjacent to one another. In order to be considered successful, the subject must locate the hidden food item without searching in the location that was not manipulated.

#### Rotation (9 trials)

In the third task, we examined subjects’ ability to track a food reward as it is spatially rotated either 180 or 360 degrees. In this task, subjects watched as one of three possible locations was baited and then as the three locations were rotated as a unit on a horizontal plane. Three different manipulations were employed. In 180 degree middle trials, the middle location was baited and the platform was turned 180 degrees. In 360 degree side and 180 degree side trials, both the left or right location was baited and the platform was then rotated 360 or 180 degrees, respectively. Subjects successfully completed a Rotation trial by tracking and identifying the correct location.

#### Transposition (9 trials)

In this task, subjects watch as a food reward is hidden in one of three possible locations and then as the baited location is changed in one of three ways. In one condition, the baited location is switched with one of the unbaited locations. In the second condition, the baited location is switched with one of the unbaited locations and then the two unbaited locations are switched. In the last condition, the baited location is switched with one of the unbaited locations and then with the other unbaited location. To be considered successful on this task, the subject must track the reward and choose the baited location.

#### Relative Numbers (13 trials)

In the fifth task, subjects were tested for their ability to discriminate between different quantities when being presented with two plates containing different amounts of equally sized pieces of food. Each subject received the same set of 13 different quantity pairings as those used in the original PCTB (1:0, 5:1, 6:3, 6:2, 6:4, 4:3, 3:2, 2:1, 4:1, 4:2, 5:2, 3:1, and 5:3). During each trial, the subject was allowed to choose only one plate and received whatever reward was on the chosen plate. A correct response was recorded when the subject chose the plate containing the larger quantity of food. We did not include the task by Herrmann et. al. (2007a) referred to as Addition Numbers.

#### Causality Noise (6 trials)

In the sixth task, we assessed subjects’ understanding of causal relationships based on sound. In this task, the experimenter placed a hard food reward (i.e., peanut) in one of two metal containers such that the container with the food reward made a sound when shaken while the unbaited container did not. In “Full” trials, the metal container containing the food reward was lifted and shaken and then the unbaited container was lifted. In the “Empty” trials, the empty container was lifted and shaken and then the baited container was lifted. Subjects were then allowed to choose one of the two containers. A correct choice was recorded when the subject chose the baited container.

#### Causality Visual (6 trials)

In the seventh task, subjects were tested for their causal understanding of the physical world in the visual domain. Specifically, in one trial type a food reward was placed underneath one of two boards laying flat on the testing table. The food caused the baited board to be tilted away from the subject (such that the chimpanzee could not see the food underneath the board) while the unbaited board remained flat. In the second trial type, a food reward was placed underneath one of two pieces of cloth laying flat on the testing table. The reward created a visible bump in the baited cloth while the unbaited cloth remained flat. In both trial types, the subject had to choose the baited item to be considered successful.

#### Tool Properties (6 trials)

This Physical Cognition task explored the apes’ understanding of the physical properties of tools and how those relate to achieving a goal. In each task, the subject was presented with a choice between two similar tools. However, one tool could be used to obtain a food reward while the other tool was ineffective. For the first task, subjects were presented with two identical pieces of paper. One piece of paper has a food reward sitting on top of the far end while the second piece of paper had a food reward sitting beside it. The subject could pull either piece of paper into their cage, but only by pulling the paper with the food sitting on top of it would they be able to retrieve the food reward. In the second task, one tool was identical to the effective tool in the first task. The second tool consisted of two smaller pieces of paper with a small gap between them, visually emphasizing that they were not connected. The food reward was placed on the out-of-reach piece of the two disconnected pieces of paper. The subject could pull in the reward using the effective tool but pulling the piece of the disconnected paper was ineffective in obtaining the reward. Note that we did not include three tool properties tasks from the original PCTB including “Bridge”, “Broken Wool” and “Tray Circle” (Herrmann et al., 2007a).

#### Tool Use (1 trial)

The last Physical Cognition task measured each ape’s ability to solve a simple problem using a tool. A food reward was placed on a table approximately 10 cm out of reach of the subject. A lollipop stick measuring 12 cm long was then placed on the table with one end inserted into the enclosure. To be successful, the subject had to use the stick to manipulate the reward within his or her reach within two minutes.

### Social Cognition Tasks

There were 3 tasks within the “Social Cognition” dimension of the PCTB and they are designed to assess their initiation in joint attention (IJA) abilities, their response to joint attention (RJA) cues, and their ability to use appropriate communicative modalities based on the attentional status of a human experimenter (Attentional State).

#### Initiation of Joint Attention (IJA)

These data were taken from previously reported findings from Leavens et al. (2015). Briefly, a food item was placed outside one end of the subject’s home cage by an experimenter who then left the area to start the trial. A second experimenter then entered and positioned themselves at the opposite end of the subject’s home cage. A correct response was recorded if the ape gestured toward the food item or the experimenter while alternating their gaze between the referent and experimenter. Trials lasted 60 seconds and all subjects received 4 trials separated by at least two minutes.

#### Attentional State

Four trials were administered to test a subject’s sensitivity to an experimenter’s attentional state. An experimenter placed a food reward on the ground outside the subject’s enclosure, after which a second experimenter approached the cage and altered their attentional state in one of two ways. During the facing toward trial, the experimenter’s face and body were oriented toward the reward and the subject. During the away body-facing trial, the experimenter’s body faced the subject but his/her face was turned away. Subjects were scored as successful if they used communicative signals in the modality appropriate to the experimenter’s attentional state (i.e., he/she uses a manual gesture in toward trials, or first using an attention getting signal followed by a manual gesture in away trials).

#### Gaze Following

Four trials were administered to test a subject’s ability to follow gaze (Hopkins, Keebaugh, et al., 2014). Sitting approximately 1 meter from the subject’s enclosure, the experimenter initially captured the subject’s attention and then shift their head and eyes to gaze at a point directly behind the focal subject for a period of 15 seconds. Subjects were scored as successful if they followed the gaze of the experimenter by looking behind them. For all the measures, the percentage of correct responses computed across trials was the initial dependent measure of interest.

### Data Analysis

For the cross-sectional analysis, for each task, the percentage of correct responses was calculated for each subject. Within each chimpanzee colony, we converted the performance scores to standardized *z*-scores. We then averaged the z-scores across all physical and social cognition tasks to create separate unit weighted average (UWA) performance score (called UWA-physical and UWA-social score) (Woodley et al., 2015). For the longitudinal analysis, a similar approach was used; however, because of the retirement and subsequent relocation of the chimpanzees to the NIH funded sanctuary, the testing was limited to the tasks that assessed physical cognition only. For this set of data, we included only those chimpanzees that were tested twice. Within each test period and colony, the performance scores were initially converted to standardized z-scores and then averaged across the 9 physical cognition tasks within the PCTB. Lastly, because we have previously found that (1) PCTB performance is significantly heritable in chimpanzees (Hopkins, Russell, et al., 2014) and (2) some of the chimpanzees in our sample were related to each other, we used relatedness coefficient scores as covariates in all analyses. Relatedness coefficients were created for the chimpanzee population using available pedigree information going back as early as the founder animals within each population (Hopkins et al., 2015). All data were analyzed in SPSS Statistics 26 (IBM Corporation, Chicago, IL, USA). Data are available from the corresponding author upon reasonable request.

## Results

### Cross-Sectional Findings

For this analysis, we performed stepwise multiple regression on each of the cognition scores entering the following variables: sex, linear age, and quadratic age in that order. In addition to the overall model results, the stepwise regression also reported significant changes in R-squared with the inclusion of each predictor variable. For UWA-physical *F*= 2.605; df = 3, 214; *p* = .050; *R*^*2*^ = .035 and UWA-social *F*= 5.103; df = 3, 214; *p* = .002; *R*^*2*^ = .067, the full model regression analyses were significant. For the UWA-social measure, significant changes in R-squared were found when the variables linear age *F*=9.871; df = 1,215; *p* = .002 and quadratic age were entered into the model *F*= 5.209; df = 1, 214; *p* = .023. For the UWA-physical score, significant changes in R-squared were only found when quadratic age was entered into the model *F* = 5.417; df = 1, 214; *p* = .0218. Scatterplots of the quadratic association between age and each cognition measure are shown in Figure 1a and 1b. As can be seen, younger and older apes performed more poorly than middle-aged apes.

**Figure 1.**
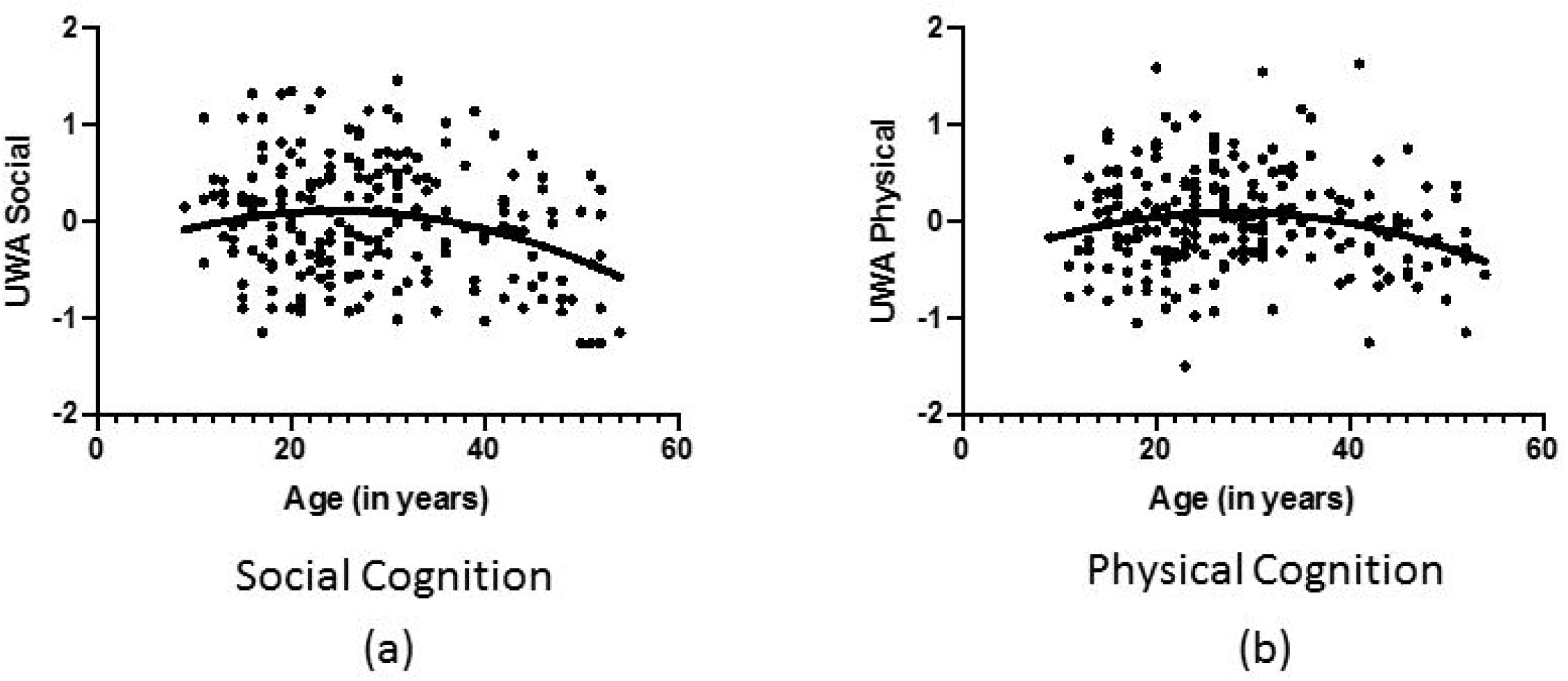
Scatterplots of quadratic association between age and (a) UWA-social and (b) UWA-physical cognition in chimpanzees.

In addition to multiple regression analyses, we also tested for age-related differences in the UWA-physical and UWA-social scores using a mixed model analysis of covariance on age groups. Performance test (UWA-physical, UWA-social) was the repeated measure while sex (male, female) and age (young, middle, elderly) group were the between group factors. Genetic relatedness was a covariate. A significant main effect for age group was found *F*=14.312; df = 1, 211; *p* = .000. Post-hoc analysis indicated that elderly chimpanzees performed worse than middle-aged and young chimpanzees but there was no difference between these two latter groups (see Figure 2a).

**Figure 2.**
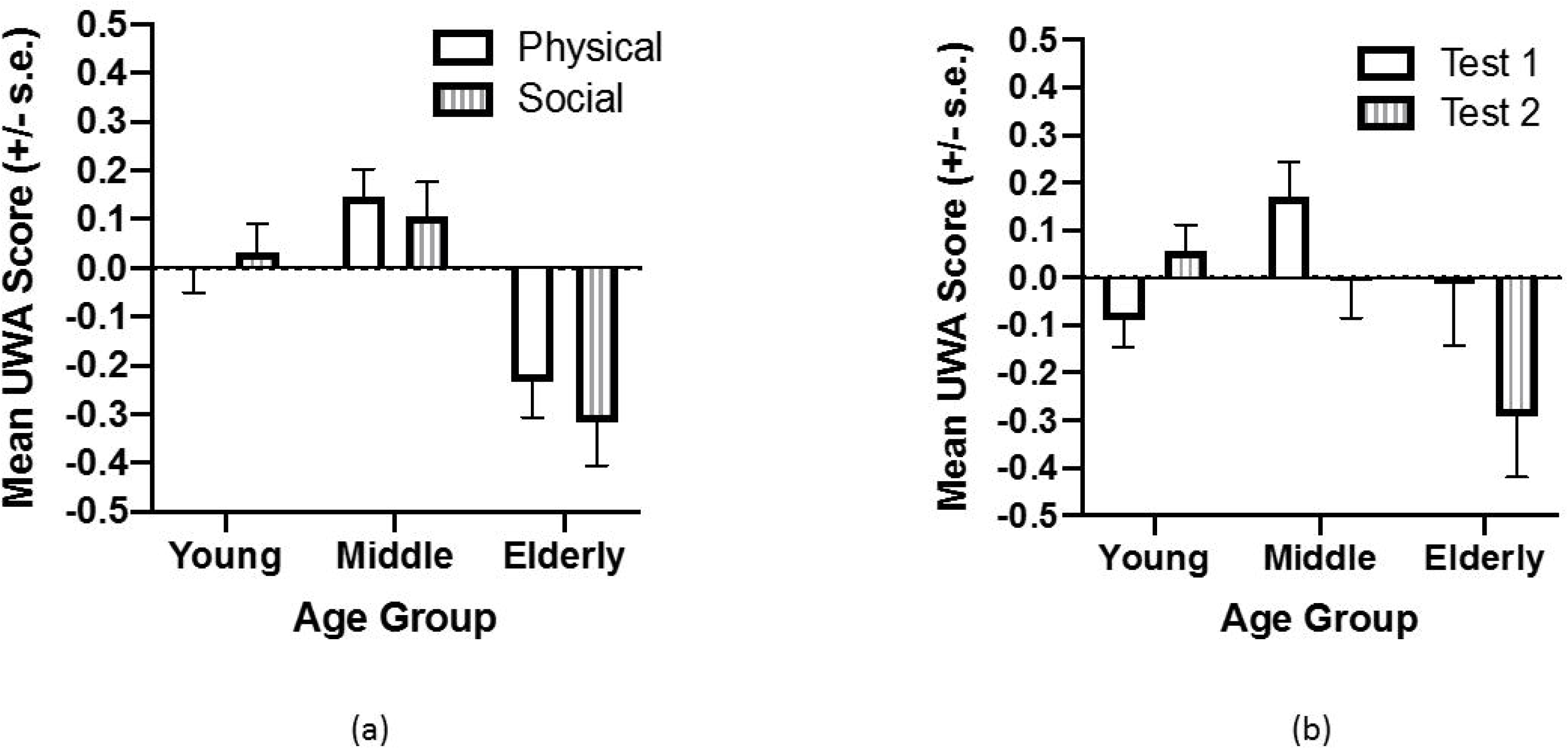
(a) Mean UWA-physical and UWA-social scores (+/− SEM) for young, middle-aged and elderly chimpanzees for the cross-sectional analysis (b) Mean UWA scores for test 1 and 2 (+/− SEM) for young, middle-aged and elderly chimpanzees in the longitudinal analysis. Note that the UWA scores reflect standardized scores and, as such, values at or close to zero reflect the group average performance. Values increasing above and below zero indicate better or worse performance relative to the average for the group.

### Longitudinal Findings

For this analysis, we performed a repeated measures analysis of covariance with test session (UWA-physical score from test 1 and test 2) as the repeated measure while age group and sex were the between group factors. Covariates included genetic relatedness and the difference in age between test 1 and 2. We found a significant two-way interaction between age group and test, *F*= 6.571; df = 2,134; *p* = .002 (see Figure 2b). Post hoc analysis indicated that, within test 1, middle-aged chimpanzees performed better than younger but not older individuals. In contrast, for test 2, the elderly chimpanzees performed worse than both the middle-aged and young individuals. As can be seen in Figure 2b, whereas performance by the chimpanzees in the young group improved from test 1 to test 2, performance between the two test periods decreased in both the middle-aged and particularly the elderly chimpanzees. No other significant main effects or interactions were found.

## Discussion

As has been reported in other nonhuman primate species using cross-sectional methods (Hara et al., 2012; Herndon et al., 1997; Joly et al., 2014; King & Michels, 1989; LaClair & Lacreuse, 2016; Lacreuse et al., 1999; Lyons et al., 2004; Moss et al., 1988; Munger et al., 2017; Nagahara et al., 2010; Picq, 2007; Rapp & Amaral, 1989, 1992; Workman et al., 2019), we found an inverted U-shaped pattern of lifespan change in cognition, where middle-aged individuals performed best on both physical and social cognition tasks compared to younger and elderly chimpanzees. The longitudinal analysis also showed age-related decline in cognition in chimpanzees, particularly those classified as elderly during the initial test. Longitudinal studies of age-related changes in cognition are rare in nonhuman primates (Workman et al., 2019) and, in the only other longitudinal study in chimpanzees, Lacreuse et al. (2014) reported a small but non-significant loss in cognition in a sample of 43 female chimpanzees, a finding that differs somewhat from the results reported here. The differences in findings between Lacreuse et al. (2014) and this report may be attributed to the larger sample size and the inclusion of male chimpanzees in the current study.

There are at least two limitations to this study. First, though the collective tasks within the PCTB did detect cross-sectional and longitudinal changes in cognition, it is possible that other kinds of tasks, such as those assessing memory and executive functions, may capture even more robust changes in cognition associated with age in chimpanzees. Second, the length of time between assessments were (1) relatively short, (2) limited to the physical cognition tasks, and (3) not well controlled or systematically manipulated in this sample. Though we used the duration in time that elapsed between the two tests as a covariate in the statistical analysis, ideally this should be better controlled from an experimental standpoint. This could have removed some potential sources of error variance and enhanced the rigor of the design. Additionally, given the longer lifespan of chimpanzees compared to other nonhuman primates, the elapsed time between tests should perhaps be longer which may reveal more robust effects of age on cognition.

There is one fundamental question that arises from these findings when viewed in the context of the neurobiology of aging in chimpanzees. Notably, chimpanzees show age-related changes in gray volume and white matter integrity (Autrey et al., 2014; Chen et al., 2013), and a proportion of elderly individuals develop AD pathology (Edler et al., in press) or exhibit other neurological pathologies or infarcts (Hopkins & Latzman, 2017). Whether the loss in cognitive functions reported here in elderly chimpanzees are attributable to any of these conditions, but specifically those pathological features associated with AD, remains entirely unknown, yet warrants further investigation. If loss in cognition in aged chimpanzees is associated with AD pathology, this would have significant implications for our understanding of the uniqueness of neurodegenerative disorders in humans compared to other nonhuman primates. Additionally, such findings would have significant implications for the long-term management and care of the captive chimpanzee populations in the US and other countries. For instance, the National Institutes of Health owns more than 400 chimpanzees, most of whom are retired from biomedical research and housed in the federal sanctuary (i.e., Chimp Haven). Many of these chimpanzees are adults, will continue to age and may potentially develop motor and cognitive impairments that present management challenges or compromises their wellbeing. In our view, this is a remarkable circumstance and opportunity in which continued documentation of longitudinal changes in cognition and behavior could be coupled with a rigorous, sustained brain collection (www.chimpanzeebrain.org) and a neuropathology program to produce data that benefit both our understanding of AD pathology in humans and the opportunity to advance wellbeing in captive chimpanzees.

One intriguing alternative possibility is that the cognitive loss we observe in chimpanzees may be relatively mild and due to subtle neuroanatomical changes in dendritic and synaptic function, but not linked with overt human-like AD pathology, despite a large number of chimpanzees exhibiting AD-like pathology on postmortem examination (Edler et al., 2017). This would be consistent with the fact that chimpanzees with AD pathology do not exhibit significant neuron loss (Edler et al., in press), a characteristic that is a hallmark of human AD. Some studies indicate a divergence between humans and chimpanzees in terms of the inflammatory responses associated with NFTs and amyloid plaques (Edler et al., 2018; Munger et al., 2018), which may contribute to neuron loss in humans and neuron preservation in chimpanzees.

In summary, the results reported here demonstrate that chimpanzees show age-related changes in cognition. This is, by far, the single largest cross-sectional and longitudinal study on age-related changes in cognition in chimpanzees, the species most closely related to humans and the only known one to naturally develop AD pathology similar to humans. Taken together, these lines of evidence highlight the importance of including chimpanzees in future research that may provide important clues that will allow us to identify therapeutic targets for human AD.

## Acknowledgements

This research was supported in part by NIH grants NS-42867, NS-73134, and HD-60563 The NCCC chimpanzees are supported by Cooperative Agreement U42-OD011197. The Yerkes Center and NCCC are fully accredited by the AAALAC International. American Psychological Association guidelines for the ethical treatment of animals were adhered to during all aspects of this study. The authors have no conflict of interest to declare.

